# Protein secondary structure and remote homology detection

**DOI:** 10.1101/2024.09.03.611022

**Authors:** Ali Al-Fatlawi, Md. Ballal Hossen, Ferras El-Hendi, Michael Schroeder

**Affiliations:** Biotechnology Center (BIOTEC), Center for Molecular and Cellular Bioengineering, Technische Universität Dresden, Dresden, Germany; Center for Scalable Data Analytics and Artificial Intelligence (ScaDS.AI), Dresden, Germany; University of Kufa, Iraq

## Abstract

A protein can be represented by its primary, secondary, or tertiary structure. With recent advances in AI, there is now as much tertiary as primary structural data available. Fast and accurate search methods exist for both types of data, with searches over both representations being highly precise. However, primary structure data can sometimes be incomplete. As a result, tertiary structure has become the gold standard for remote homology detection.

How does secondary structure perform in remote homology detection? Secondary structure interprets proteins as a sequence using an alphabet representing helices, strands, or loops. It shares its sequential nature with primary structure while retaining topological information similar to tertiary structure.

To assess the effectiveness of secondary structure in remote homology detection, we devised a challenging classification task aimed at determining the superfamily membership of very distantly related protein domains. We used benchmarks from the CATH and SCOP databases and evaluated sequence and structure alignment algorithms on primary, secondary, and tertiary structures.

As expected, both basic and advanced sequence alignment algorithms applied to primary structure achieved high precision, but their overall area under the curve was lower compared to the gold standard of structural alignment using tertiary structure.

Surprisingly, a simple string comparison algorithm applied to secondary structure performed close to the gold standard. This result supports the hypothesis that key structural information is already encoded in secondary structure and suggests that secondary structure may be a promising representation to use when high-confidence structural data is unavailable, such as in cases involving protein flexibility and disorder.

## 2 Introduction

The primary structure represents proteins as sequences of amino acids, while the tertiary structure provides a set of atomic coordinates of these amino acids, which form helices, strands, or loops in 3D space. The secondary structure serves as an intermediate representation, capturing helices, strands, and loops in a sequential format. Generally, tertiary structure is more conserved than primary structure, as functional requirements impose constraints on a protein’s structure, whereas the sequence itself can mutate as long as essential functions are preserved [1]. As a result, dissimilar sequences may fold into similar structures that are homologous in function [2, 3]. Functional constraints often apply only to specific regions of the protein rather than the entire structure, and a substantial portion of the protein sequence can undergo significant variation as long as the core structure, which satisfies functional requirements, remains intact [4]. Consequently, primary structure alignment tools like BLAST [5] and HHblits [6] are highly precise in remote homology detection, but they are not complete for the reasons mentioned above. On the other hand, tertiary structure serves as a gold standard, offering both precision and completeness [3].

In contrast to sequence alignments, naive structural alignments performed by tools like TM-align [7] and CE-align [8] are computationally intensive, as these algorithms determine similarity by finding an optimal superposition of atomic coordinates. Such methods are not suitable for searching large structural repositories, such as the AlphaFold database [9] or the ESM Metagenomic Atlas [10], which contain hundreds of millions of structures. Therefore, 3D fingerprints have been developed and implemented in the structure search engine Foldseek [11]. These 3D fingerprints represent proteins as sequences over an abstract alphabet, capturing local information about the atoms’ coordinates and interactions. In summary, fast and accurate searches over primary and tertiary structures are available with BLAST and FoldSeek, respectively. The former is highly precise in detecting remote homologs, while the latter is both precise and complete.

How does secondary structure perform in homology detection? Does it share the precision of BLAST but fail to retrieve all remote homologs, or does its abstract representation hold sufficient structural information to perform as well as tertiary structure? The latter hypothesis is supported by Przytycka et al., who explored the extent to which a protein’s secondary structure could inform its three-dimensional fold by analyzing known protein structures [12]. They constructed a taxonomy based solely on secondary structure, proposing a simple mechanism of protein evolution [12]. Fontana et al. and Guharoy et al. found that secondary structure is sufficiently conserved to compute alignments of protein secondary structures against a library of domain folds [13] and to identify binding motifs in protein-protein interactions, respectively [14]. A concrete example of the conservation of secondary structure is the superfamily of single-strand annealing proteins, which comprises five distantly related families, all sharing a secondary structure motif of a *β*-hairpin flanked by two helices and a *β*-sheet with a perpendicular helix [15] (see Figure 1).

**Fig. 1.**
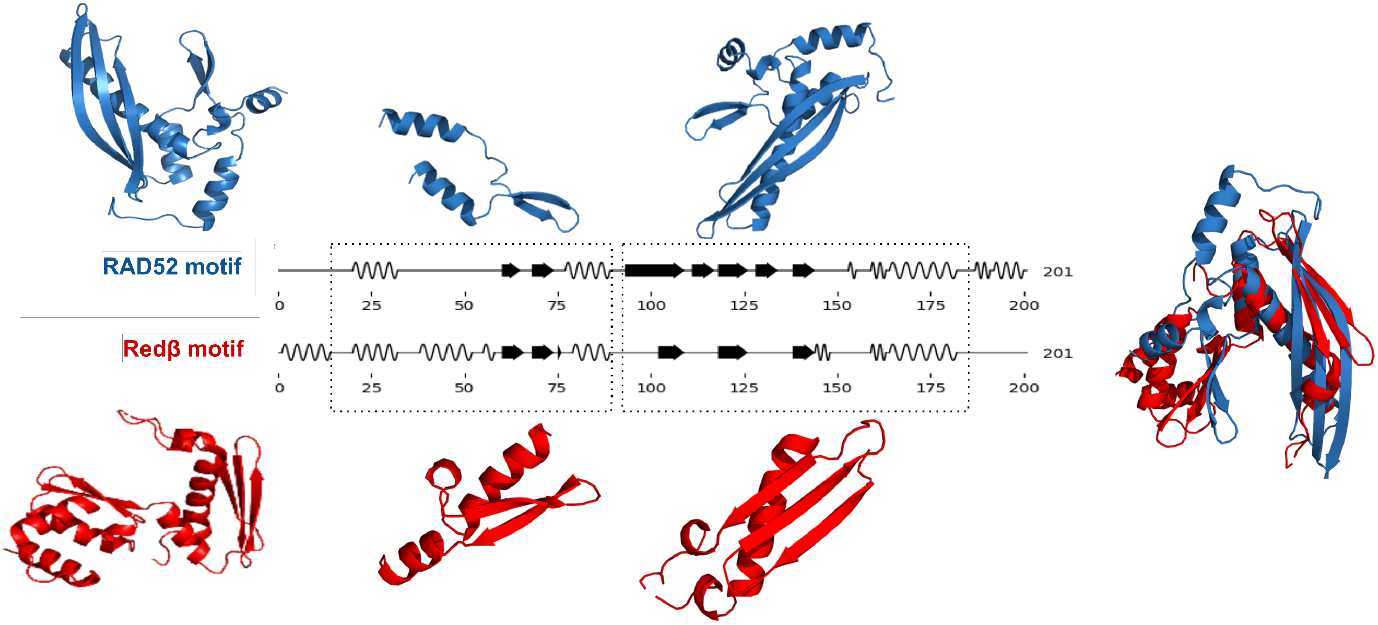
Diagram illustrating the secondary sequence alignment between Rad52 and Red*β* as an example from SSAPs.

In addition to the conservation of secondary structure, a second reason for its effectiveness is that very fast sequence alignment algorithms can be applied to secondary structure, a key factor in the fast structure search capabilities of FoldSeek using 3D fingerprints.

To answer the question of how secondary structure compares to primary and tertiary structures in remote homology detection, we focus on a challenging task: determining superfamily membership for a non-redundant set of structural domains from the CATH and SCOP domain databases. Non-redundancy refers to sequence similarity, making this task naturally difficult for sequence algorithms based on primary structure. For all three representations, we measure the similarity of domain pairs from the same superfamily versus those from different superfamilies. We aim to determine whether the performance of alignments based on secondary structure is closer to the gold standard of tertiary structure or to the highly precise but incomplete primary structure.

## 3 Results and Discussion

### Secondary structure alphabet

Secondary structure is an abstraction of 3D structure. In 3D, the connection between two consecutive amino acids is described by two angles, known as the phi and psi angles. These angles cannot adopt all theoretical combinations but instead cluster around certain values. These clusters lead to the assignment of secondary structure to a residue. Depending on the clustering, the secondary structure can be represented using a more restrictive or a less restrictive alphabet. Typically, two representations are used: a 3-letter alphabet (helix, strand, loop) and an 8-letter alphabet (which includes three types of helices 3_10_-helix, *α*-, and *π*-helix—strand, loop, and three additional letters for specialized turns, strands, and coils). We compare both representations to determine whether the increased granularity of the 8-letter alphabet leads to improved performance.

### Benchmark datasets

To compare primary, secondary, and tertiary structures, we employed a challenging classification task. We compared scores within predefined domain superfamilies and between distinct superfamilies, where the former should yield better scores. The two principal databases for structural domain classification are CATH [16] and SCOPe [17], which organize protein domains by topology at the family and superfamily levels. We included both databases and varied the degree of redundancy in our datasets. A non-redundant dataset is more challenging, especially for sequence-based methods, but also more realistic for scenarios involving remote homology detection, where little prior knowledge exists. Therefore, we devised three datasets: CATH (with redundancy), SCOPe40 (non-redundant at 40% sequence similarity), and CATH S20 (non-redundant at 20%). Each dataset covers over 1,000 superfamilies and more than 10,000 individual domains. The number of pairs ranges from 62 to 286 million, with the number of pairs within the same superfamily being a small fraction of those between different superfamilies (see Table 1), ranging from 0.36% to 2.2%. This classification task is highly unbalanced, reflecting the task of remote homology detection, where the vast majority of relationships are negative. The ratio of positive to all pairs in the two non-redundant datasets, SCOPe40 and CATH S20, is an order of magnitude smaller than in the redundant dataset, CATH, highlighting the increased difficulty of these datasets for the algorithms.

**Table 1.**
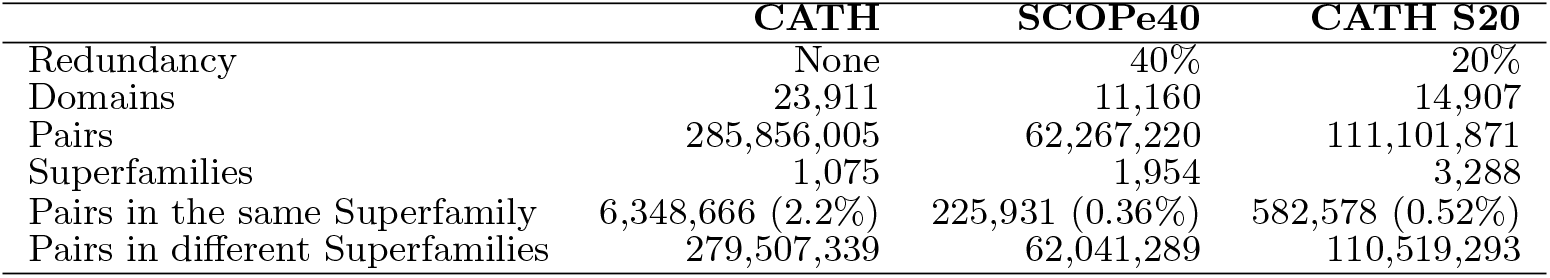
Overview of CATH and SCOPe40 datasets, including redundancy
levels and pair counts.

|  | CATH | SCOPe40 | CATH S20 |
| --- | --- | --- | --- |
| Redundancy | None | 40% | 20% |
| Domains | 23,911 | 11,160 | 14,907 |
| Pairs | 285,856,005 | 62,267,220 | 111,101,871 |
| Superfamilies | 1,075 | 1,954 | 3,288 |
| Pairs in the same Superfamily | 6,348,666 (2.2%) | 225,931 (0.36%) | 582,578 (0.52%) |
| Pairs in different Superfamilies | 279,507,339 | 62,041,289 | 110,519,293 |

### Algorithms

The Levenshtein distance [18] is the most basic sequence comparison algorithm, computing the minimal number of insertions or deletions necessary to convert one sequence into another. For comparing secondary structure sequences, we used the Levenshtein distance with one modification: we normalized it to account for substantial variations in sequence length (see Methods). The Levenshtein distance also forms the basis for advanced algorithms like BLAST, which uses an enhanced scoring scheme for gaps and mismatches and is optimized for speed. While BLAST is the standard method for amino acid sequence comparison, specialized approaches optimized for remote homology detection exist, such as those using hidden Markov models (HMMs). HMMs generate a statistical representation of a protein family, which is more robust than individual sequences. A widely used HMM implementation is HHblits [6]. For tertiary structure, the most widely used comparison method translates 3D structures into 3D fingerprints, sequences that can be interpreted as high-dimensional vectors, allowing for very efficient comparison methods. An example of such an approach is Foldseek [19]. However, since these methods aim to approximate the slower, optimal superposition of atomic coordinates, we used TM-align [20] as a reference and gold standard.

In summary, we compared primary structure using basic BLAST and advanced HHblits methods, secondary structure using a normalized Levenshtein distance with 3-letter and 8-letter alphabets, and tertiary structure using TM-align. These were applied to the three benchmark datasets: CATH, SCOPe40, and CATH S20.

### Secondary structure’s performance is closer to tertiary’s than to primary’s

Primary, secondary, and tertiary structures achieved AUCs of up to 84%, 95%, and 98%, respectively, across the three benchmarks and varying setups (see Table 2 and Figure 2). This indicates that a basic sequence alignment algorithm, such as the Levenshtein distance applied to secondary structure, significantly outperforms both basic and advanced algorithms on primary structure and approaches the performance of tertiary structure. This suggests that the topological information embedded in secondary structure can be effectively utilized, even with simple alignment algorithms.

**Table 2.** Performance (AUCs) comparison of different methods on CATH,SCOPe40, and CATH S20 datasets.

| Structure | Methods | CATH | SCOPe40 | CATH S20 |
| --- | --- | --- | --- | --- |
| Primary (amino acid) | BLAST | 0.66 | 0.56 | 0.51 |
|  | HHblits | 0.84 | 0.77 | 0.58 |
| Secondary (SS string) | Levenshtein with 3-letter | 0.95 | 0.89 | 0.81 |
|  | Levenshtein with 8-letter | 0.95 | 0.90 | 0.81 |
| Tertiary (3D) | TM-score | 0.98 | 0.95 | 0.90 |

**Fig. 2.**
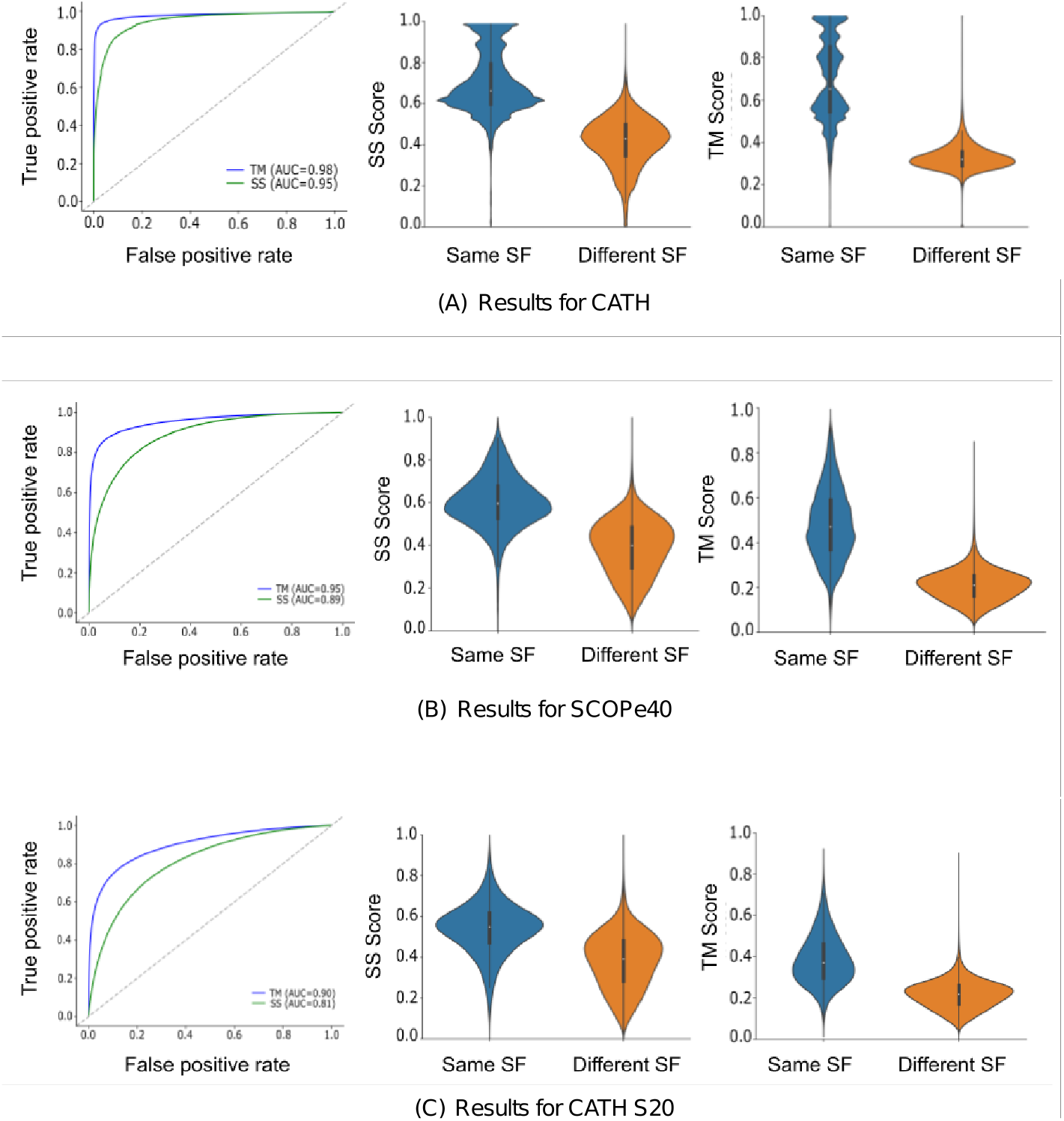
This figure presents the true positive rate (TPR) against the false positive rate (FPR), along with their corresponding AUC values, for (A) CATH, (B) SCOPe40, and (C) CATH S20. The AUC plot compares the performance of tertiary structure alignment with TM-align and secondary structure with Levenshtein using sequences in its 3-letter representation. “SS scores” refer to secondary structure alignment scores, and “SF” pertains to the superfamily. The violin plots show how the secondary structure approximates the information of the tertiary structure in separating proteins that belong to the same superfamily (Same SF) and those in different superfamilies (different SF).

### Non-redundant datasets are consistently more challenging

The more non-redundant a dataset is, the fewer “easy” remote homologues exist, making the dataset more challenging. Surprisingly, this is true not only for algorithms that explicitly leverage redundancy, such as HHblits, but also for secondary and tertiary structure-based methods.

### Performance of HMMs increases with the amount of sequences

HMMs capture a statistical signature of a sequence family, making them more effective for remote homology detection than basic sequence comparison. As a result, HMMs consistently outperform BLAST, especially as more sequence data becomes available. For instance, HMMs achieved 84% AUC on the CATH dataset, compared to 51% for BLAST on the highly redundant CATH S20. This demonstrates the effectiveness of HMMs in leveraging large sequence datasets. The recent success of embeddings computed from large language models for amino acid sequences [21, 22] builds on this effect.

### TM-Score is a gold standard

Structural alignment of tertiary structures across the full CATH dataset nearly perfectly classifies (98%) domain pairs as remote homologues or not. However, its performance drops by 8% with reduced redundancy. One reason for this is that remote homologues can vary in structure, and structural flexibility can affect alignment accuracy.

### The Secondary structure alphabet does not affect classification

One might assume that a more detailed representation of secondary structure would lead to improved results, but this is not the case. The AUC results are nearly identical for both the 3-letter and 8-letter representations. This is because, unlike amino acid alphabets, where large differences in frequencies can be attributed to physicochemical properties (e.g., the rarity of cysteines forming disulfide bonds), such specific roles are not ascribed to the more detailed secondary structure descriptions of the 8-letter alphabet (see Supplementary Material, Note 1).

### Secondary structure performs nearly as well as the gold standard

On the CATH dataset, secondary structure achieved an AUC only 3% lower than the tertiary structure gold standard. For the most difficult benchmark, this margin increases to 9%. However, compared to HMMs, which perform 32% worse than the gold standard, this is a remarkable result.

### Substitution matrices with local alignment have potential

To explore whether the 9% gap can be further reduced, we turned to more advanced sequence comparison algorithms. The Levenshtein algorithm does not employ scoring with varying gap penalties or consider likely versus unlikely mismatches. However, even though secondary structure lacks the fine granularity of primary structure, differences exist—for example, helix residues are more likely to be replaced by loop residues than by strand residues. This was observed in [23], where a scoring scheme for secondary structure was developed. To this end, we created two substitution matrices for secondary structure, inspired by BLOSUM [24]. Using high-quality multiple sequence alignments from PFAM [25], we mapped the amino acids to the corresponding secondary structure letters and computed substitution frequencies. The resulting secondary structure substitution matrix was used in a local alignment with the Smith-Waterman algorithm. On the challenging CATH S20 dataset, this approach improved performance by 5%, from 80% to 85%, narrowing the gap to the gold standard from 9% to 5% (see Supplementary Material, Note 2 and Note 3).

### Setting a threshold for secondary structure

In this study, we focused on comparing representations for which AUC is an adequate measure. However, to make practical use of secondary structure in homology detection, it is crucial to assess the likelihood of a score and establish a threshold. We utilized Bayes’ Theorem to calculate the posterior probabilities of protein pairs belonging to the same superfamily based on their secondary structure alignment scores, providing a probabilistic framework for interpreting these scores within the context of our dataset [26] (see Methods). The groups exhibited different secondary structure score ranges, with some overlap between 0.5 and 0.7. Protein pairs within the same superfamily generally had higher alignment scores, predominantly within the 0.6 to 1.0 range. For further details, see Supplementary Material, Note 4.

## 4 Methods

### Data collection

CATH version 4.3.0 and SCOPe version 2.07 were used. To maintain a fair comparison, we kept proteins with lengths between 50 and 250, representing most of the domains. To narrow our dataset further, we only considered domains with a single selection range and had the correct selection range of residues in CATH with superfamily classification. This resulted in a benchmark of 23,911 domains. The non-redundant sets of CATH S20 and SCOPe40 were fully considered without filtering; see Table 1. Secondary structures for these domains were extracted using Pymol (v 2.2.0 Open-Source) for the C*α* atoms only.

### Substitution matrix for secondary structure

For the development of the substitution matrix, Pfam-A Seed alignments were used (downloaded 23/04/24) [25]. These were filtered to only include sequences for which AlphaFold structures existed and the alignment ranges matched. This resulted in 19,226 alignments of a total of 1,155,996 sequences. Secondary structures for these sequences were extracted again as above. In accordance with the original BLOSUM authors [24], the alignments were then trimmed to remove columns containing gaps, leaving 2,460,186 ungapped columns.

With the data prepared, we apply the same steps as for BLOSUM. Firstly, pairwise frequencies *f*_*ij*_ are counted for all columns and all pairs *ij*. Next, the observed probability *q*_*ij*_ is calculated as the frequency of a pair *ij*, relative to the number of all pairs:

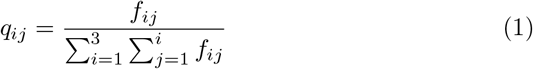

Next we calculate the probability of occurrence *p*_*i*_ for all possible letters *i* as

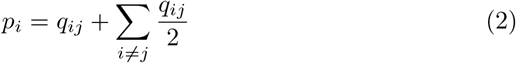

With this we can determine the expected probability *e*_*ij*_ with *e*_*ij*_ = *p*_*i*_*p*_*j*_ for *i* = *j* and 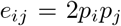 for *i j*. A log-odds ratio is then calculated as *s*_*ij*_ = log_2_(*q*_*ij*_*/e*_*ij*_ and lastly, these scores are then rounded to the nearest integer and multiplied by 2 to determine the final substitution matrix values.

### Sequence and structural alignments

The Levenshtein algorithm [18] aligns secondary structure sequences using the Levenshtein Python C extension module (v 0.20.9). To compute the scores, we divided the Levenshtein distance by the maximum length (number of residues) of the two sequences and subtracted this value from one.

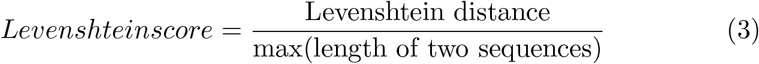

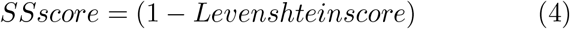

For local alignment, we implemented the Smith-Waterman algorithm [27], incorporating a custom substitution matrix as described above and affine gap penalties akin to the one used in BLAST [28]. The gap opening penalty was set to -11, and the gap extension penalty was set to -3. Finally, we computed the alignment scores (SS score) by dividing the similarity score by the longer of the two sequences’ maximum score (score if compared to itself). For the structural alignment, we used US-Align [29], a successor of the TM-align algorithm [20].

### Visualization

PyMol (v 2.5.0) was used to visualize the structures. The Biotite python package (v 0.37.0) has been used to visualize the secondary structure aligned strings [30]. Seaborn (v 0.12.2) was used to generate violin plots and other types of plots.

### Posterior Probabilities

We followed the workflow of [31] for calculating posterior probabilities. The posterior probability of a pair of proteins belonging to the same superfamily at a certain SS-score can be calculated using Bayes’ Theorem:

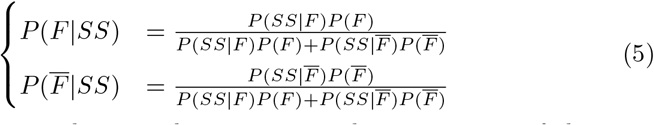

Here, SS represents the secondary structure alignment score of the compared proteins as calculated by our method SS score *F* and 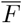
represents the case that the proteins belong to the same and different CATH superfamily, respectively; *P* (*F*) and 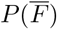are the prior probabilities of proteins in the same and different superfamily;
*P* (*F* |*SS*)and 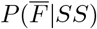 are the conditional probabilities of SS score when the two proteins are in the same or different superfamily, respectively. The number of occurrences in our dataset can determine these conditional probabilities:

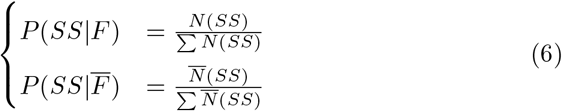

where *N* (*SS*) is the number of protein pairs with a certain SS score (SS) and 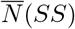 is the number of protein pairs with the SS score in the different superfamily. The denominators are the summation of the same and different superfamily protein pairs for all SS scores in (0, 1], which equals the total number of protein pairs in our dataset, respectively. To calculate the *a priori* probabilities, we used the all-to-all pairing on our dataset. The prior probabilities can then be calculated by

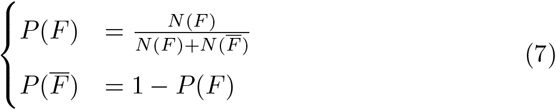

where *N* (*F*) and 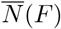 are the numbers of the same superfamily and the different superfamily pairs. With all these preparation steps, we can now calculate the *a posteriori* probabilities according to Equation 5.

## 5 Conclusion

The representation of proteins is an intriguing and open-ended question, with the answer depending on the specific purpose and requirements of the representation. Until recently, there was a trade-off: representing proteins as primary structures allowed for a wealth of data and fast algorithms, which was not the case for tertiary structures. However, with the advent of AlphaFold and related systems, there is now a comparable amount of structural data, and with the development of FoldSeek, structural representations are also amenable to fast structural searches. As a result, fast and accurate remote homology detection is now possible, leveraging tertiary rather than primary structures.

While this question seems practically settled, one aspect remains open: how does secondary structure perform in remote homology detection? Secondary structure shares the sequential nature of primary structure and the topological information of tertiary structure. In this work, we address this question and report the surprising result that simple sequence comparison of three-letter secondary structure performs nearly as well as tertiary structure in distinguishing domains from the same superfamily. Moreover, secondary structure offers an advantage that tertiary structure lacks: flexibility in tertiary structure can make similarity detection difficult for structure alignment algorithms, whereas secondary structure is robust against such variations. Therefore, secondary structure is a promising representation that should be considered when dealing with 3D protein structures.

## 6 Data Availability

The secondary structure strings, SS scores, TM scores, and superfamily memberships are available at the following link:

https://sharing.biotec.tu-dresden.de/index.php/s/eKaamoJtTJPLbJk

## 7 Acknowledgement

We gratefully acknowledge the financial support provided by the BMBF projects scads.ai and SNRT, as well as the access to high-performance computing resources through the ZIH of TU Dresden, which were instrumental in the completion of this study. We extend our sincere thanks to Francis Stewart and Christian Pilarsky for their valuable discussions and insights, which greatly contributed to the development of this research.

## 8 Author contributions

AAF and MS conceived the study. AAF, BH, FE, and MS analyzed data and wrote the manuscript.

## Supplementary Material

**Supplementary Note 1:**

**Fig. S1.**
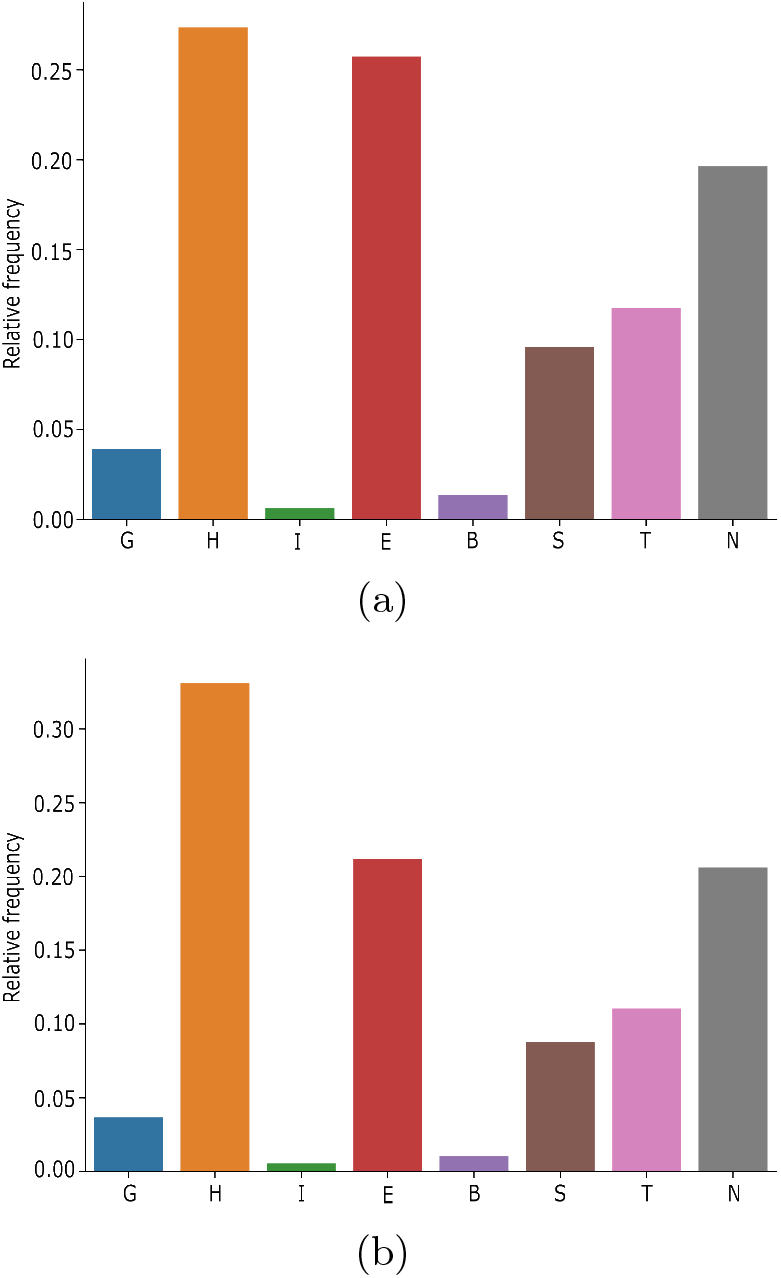
The frequency of each letter in the 8-letter representation secondary structure according to the DSSP method. (a) CATH dataset (b) SCOPe40

**Supplementary Note 2:**

***The likelihood of replacement for three secondary structure letters (S, H, L) in CATH and SCOPe40***

**Fig. S2.**
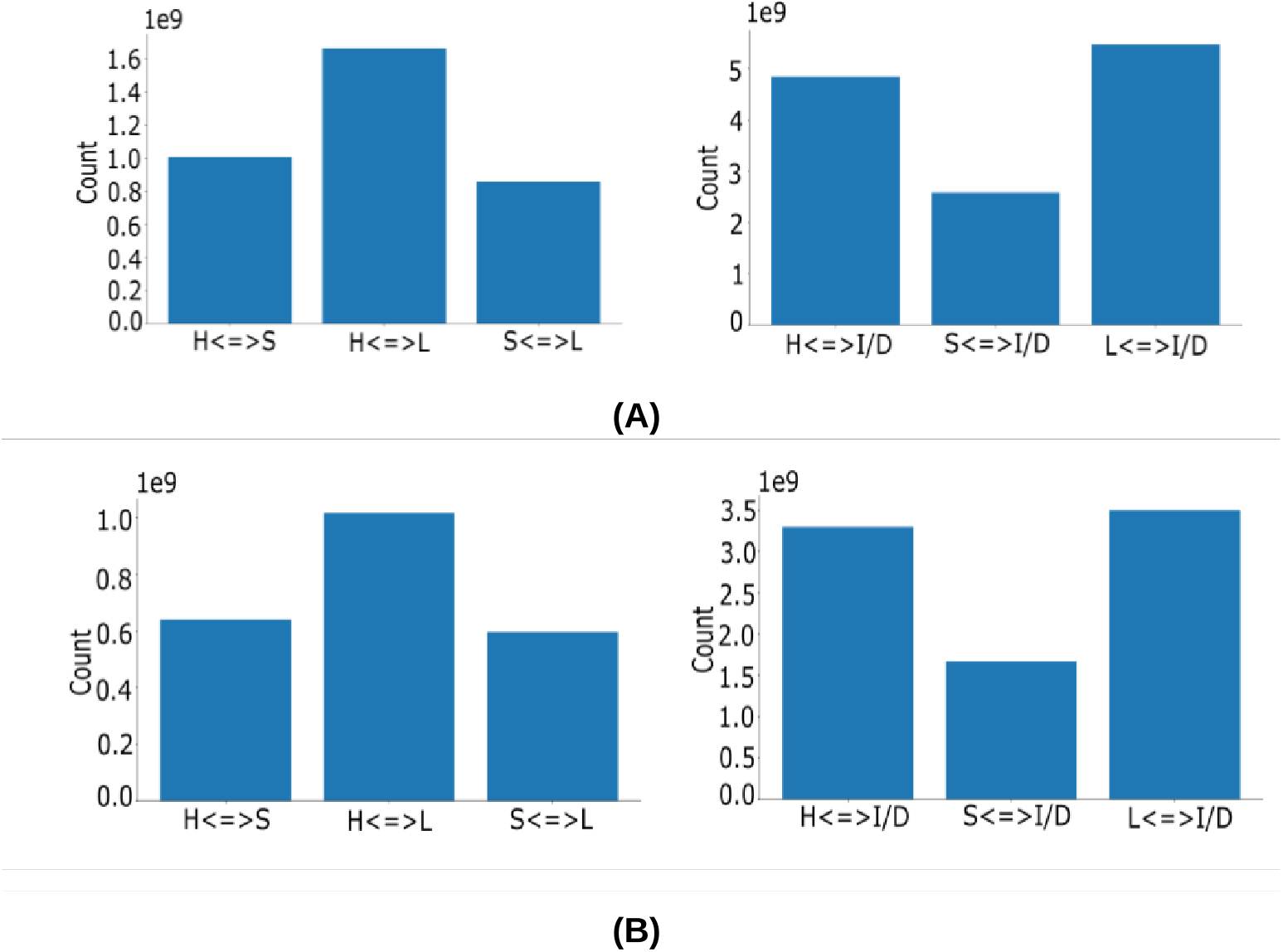
Substitution counts for letters in CATH (A) and SCOPe40 (B) datasets. ‘H’ represents helices, ‘S’ represents sheets, ‘L’ represents loops, and ‘I/D’ stands for insertion/deletion events. The ¡=¿ symbol indicates bidirectional substitution. In both panels (A) and (B), the left figures show substitution counts between ‘H’, ‘S’, and ‘L’, while the right figures display substitution counts among ‘H’, ‘S’, ‘L’, and insertions/deletions.

**Supplementary Note 3:**

***Custom substitution matrix for secondary structure derived using log-odds ratios from Pfam-A Seed alignments***

**Table S1.** Pairwise substitution values used in the Smith-Waterman alignment algorithm for SS strings.

|  | H | S | L |
| --- | --- | --- | --- |
| H | 2 |  |  |
| S | -7 | 3 |  |
| L | -16 | -5 | 4 |

**Table S2.**
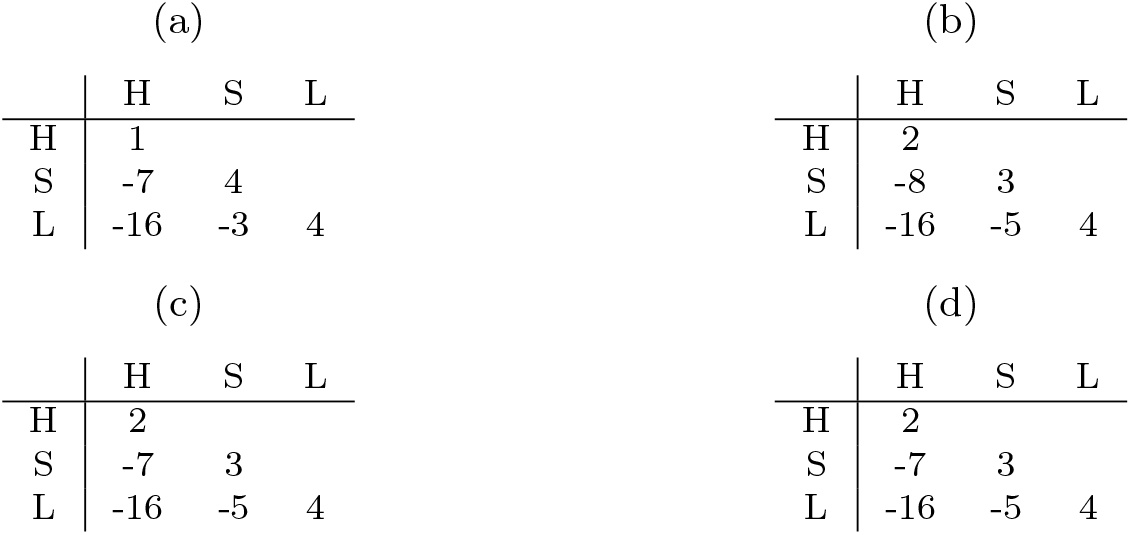
In order to verify the robustness of the matrix, we re-calculated the matrix for smaller sample sizes, of **a)** 500, **b)** 1,000, **c)** 5,000 and **d)** 10,000 alignments, that were randomly sampled from the full Pfam-A Seed alignment set. As can be seen from a) and b) for smaller sample sizes, there are minor deviations of magnitude up to 2 in a) and 1 in b) respectively, however for a sample size of 5,000 the substitution matrix values match the default matrix’s values, such that it is safe to assume our matrix to be robust. We ran a similar experiment to test robustness against sequence similarity, where matrices were computed using only alignments of maximum pairwise similarity of 20%, 45%, 50%, 62%, 80%, and 90% respectively. All of these matched the original matrix, which is not surprising since domains are considered to have structurally conserved motifs.

**Fig. S3.**
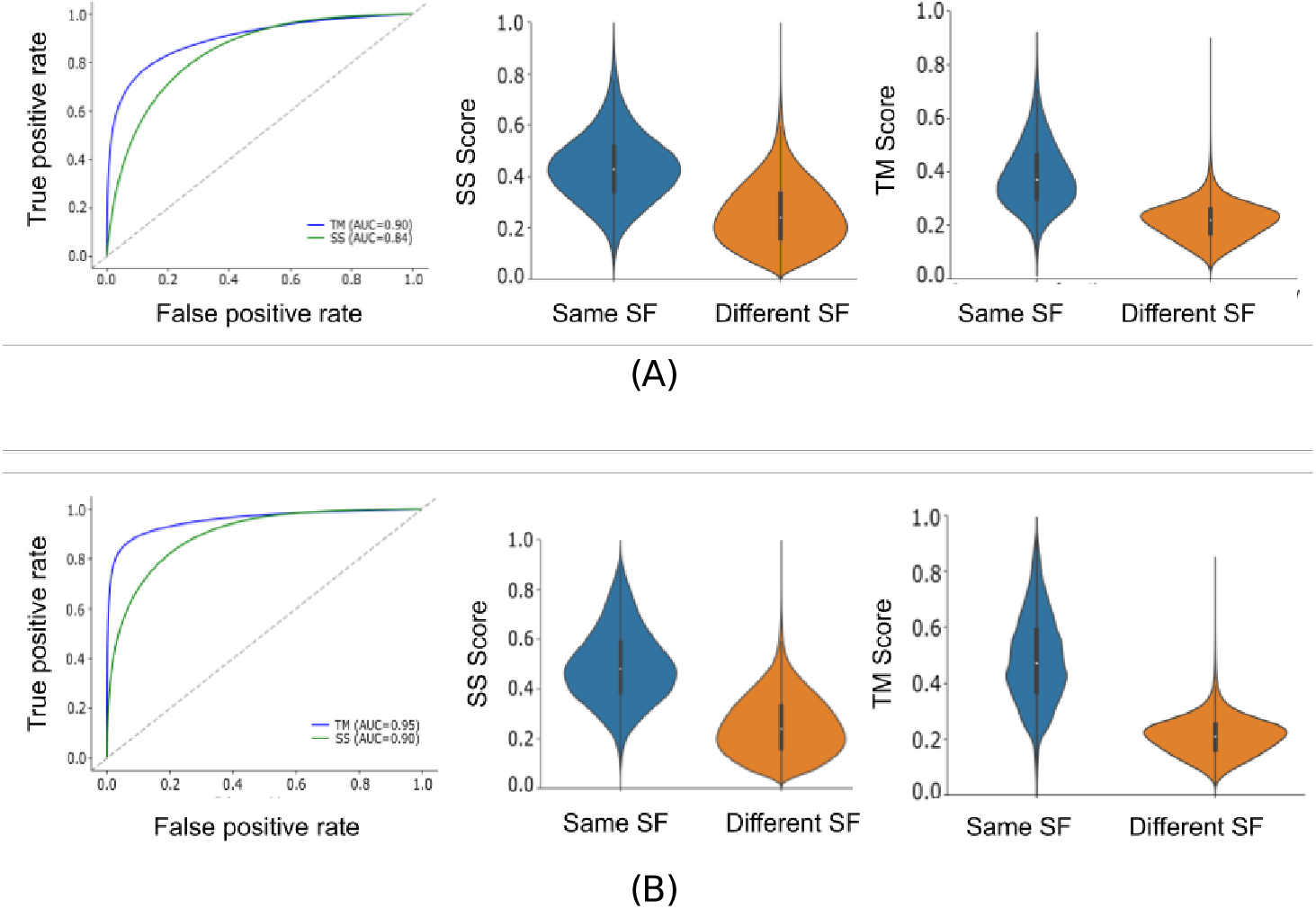
True Positive Rate (TPR) vs. False Positive Rate (FPR) and corresponding AUC values for CATH S20 (A) and SCOPe40 (B), using a local alignment with a customized substitution matrix. We set the penalty for replacing ‘S’ with ‘L’ or ‘H’ with ‘L’ (i.e., -0.5) lower than for replacing ‘S’ with ‘H’ or vice versa, which was set at -1 (see Methods). This customization was driven by the observed distribution of replacements between these letters. Here, we used an alternative Needleman-Wunsch algorithm [15] for local alignment. These modifications improved performance in CATH from 81% to 84% and slightly enhanced performance in SCOPe from 89% to 90%

**Supplementary Note 4:**

***Further Statistical Analysis***

We calculate the conditional probability for a given secondary structure alignment score (SS score) of protein pairs being in the same superfamily. For an SS score below 0.6, the probability of a protein pair belonging to the same superfamily is close to 0; this is the case for only a few pairs. However, we can also observe that for an SS score above 0.8, that probability increases sharply to around 50%. Around the score of 0.85, we observe a clear phase transition. There are also a few unexpected spikes or respective drops, especially for SS scores around 1.0, but these are rather outliers and can be explained by anomalies in and the reduced size of the dataset. Nonetheless, this phase transition points at a possible threshold of 0.85, indicating that two proteins can be expected to be in the same superfamily or superfamilies. The same procedure is done on the TM score in our data, and as was underlined in previous studies [31], the threshold of 0.5 is optimal for superfamily classification, figure below. We also performed the same analysis on the SCOPe dataset and for several methods, i.e., TM score and SS score using a 3-letter secondary structure assignment, consisting of a three-letter alphabet, and the DSSP secondary structure assignment, consisting of an eight-letter alphabet [32, 33].

**Fig. S4.**
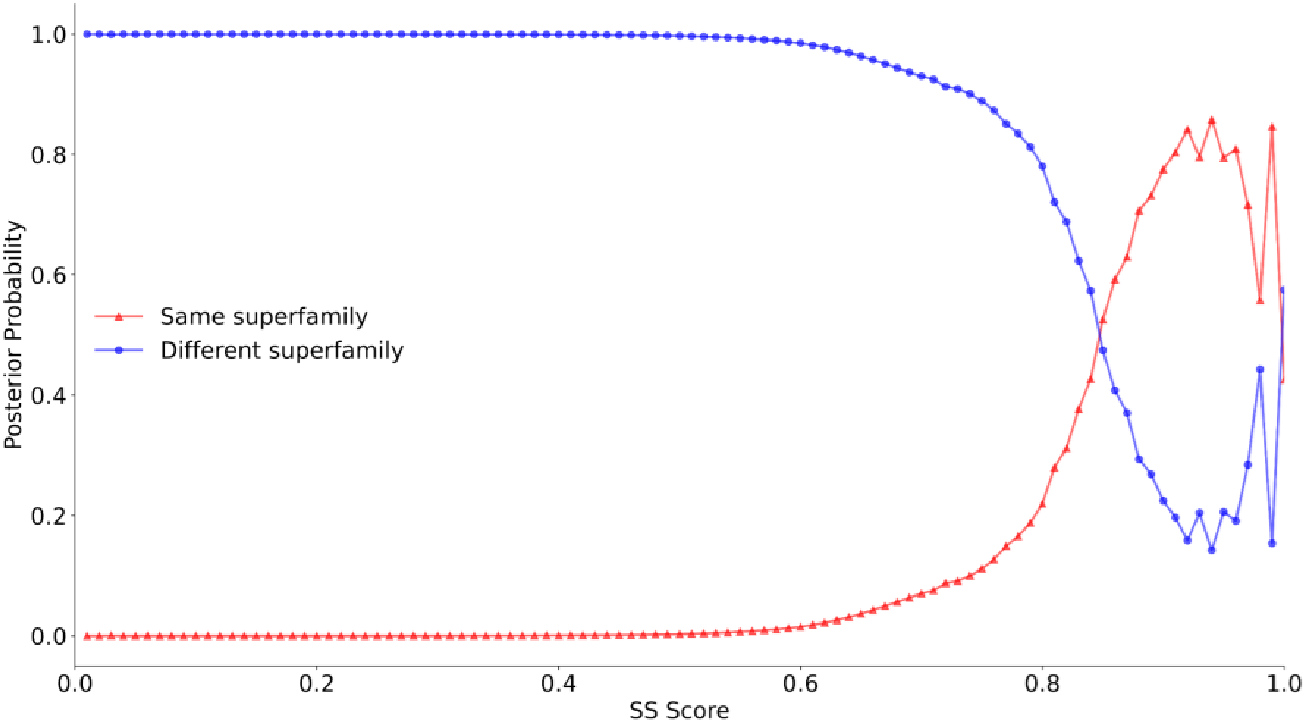
The posterior probabilities of SCOPe domain pairs for a given secondary structure score being in the same superfamily (red) or different superfamilies (blue). Both lines cross at around 0.85.

